# Exploring Drivers of Gene Expression in The Cancer Genome Atlas

**DOI:** 10.1101/227926

**Authors:** Andrea Rau, Michael Flister, Hallgeir Rui, Paul L. Auer

## Abstract

The Cancer Genome Atlas (TCGA) has greatly advanced cancer research by generating, curating, and publicly releasing deeply measured molecular data from thousands of tumor samples. In particular, gene expression measures, both within and across cancer types, have been used to determine the genes and proteins that are active in tumor cells. To more thoroughly investigate the behavior of gene expression in TCGA tumor samples, we introduce a statistical framework for partitioning the variation in gene expression due to a variety of molecular variables including somatic mutations, transcription factors (TFs), microRNAs, copy number alternations, methylation, and germ-line genetic variation. As proof-of-principle, we identify and validate specific TFs that influence the expression of *PTPN14* in breast cancer cells. We provide a freely available, user-friendly, browseable interactive web-based application for exploring the results of our transcriptome-wide analyses across 17 different cancers in TCGA at http://ls-shiny-prod.uwm.edu/edge_in_tcga.

## Introduction

Large-scale genomics projects, such as The Cancer Genome Atlas (TCGA), have greatly advanced biomedical research by generating, curating, and publicly releasing multiple omics datasets collected from thousands of samples^2,13,19,24^. Beyond facilitating specific, hypothesis driven research, data from TCGA offer the unprecedented opportunity to conduct genome-wide integrative analyses that may offer insights not available from more directed “look-ups” of the data.

In this spirit, Jiang *et al*. (2015)^12^ integrated transcription factor (TF) profiles collected from the ENCODE project with tumor data from 6,576 TCGA samples in 17 different cancer types. They found that tumor-specific gene expression is highly influenced by TF regulatory potential after controlling for local genomic factors such as promotor methylation and nearby copy-number alterations (CNAs). In addition to the clear importance of TF expression in regulating gene expression in cancer cells, there is compelling evidence that germ-line genetic variation may exert large effects on gene expression and splicing^18^, in addition to the role of promoter methylation^21^, microRNA expression^26^, copy number alterations, and somatic changes^4^ in tumor cells.

In this work, we consider an orthogonal approach to that of Jiang *et al*. (2015) to analyze gene expression in TCGA data. Rather than focusing on how different molecular states (e.g., methylation, CNA) influence genome-wide gene expression within a subject, we instead explore the molecular drivers of per-gene expression variation across subjects in several cancer types. Our goal is thus to leverage the richness of matched multi-omic TCGA tumor data on genetic variation, methylation, microRNA expression, TF expression, copy number alterations, and somatic mutations to simultaneously estimate the relative contribution of each component on gene expression.

Structurally, our approach is similar to that of Hoffman *et al*. (2016)^11^, who analyzed the GEUVADIS data^14^ to identify different factors (e.g., sex, lab, ancestry) that influence gene expression across hundreds of subjects. We extended this approach to partition gene expression variance in the TCGA data with the goal of identifying the relative importance of molecular drivers of gene expression within cancer types. To do so, for each of 17 cancer types we considered every sample in TCGA that was assayed for gene expression, methylation, copy number alterations, somatic mutations, microRNA expression, and germ-line genetic variation. Using a linear mixed model, we partitioned the variance in gene expression (for each gene) due to these different sources. Our analyses thus provide a way to compare the relative effects of these genetic and epigenetic drivers of gene expression for each gene and cancer type.

In order to facilitate rapid exploration of the drivers of gene expression in TCGA, we have developed and deployed a free, publicly available web-based R/Shiny application called *Exploring Drivers of Gene Expression (EDGE) in TCGA* for visualizing the results of our analyses. Though the intent of our work is to provide an exploratory tool for prioritizing the drivers of gene expression within a given cancer type, we provide strong proof-of-principle of our results by validation of suggested findings from *EDGE in TCGA* with a promoter reporter assay in breast cancer cell lines.

## Methods

### TCGA data acquisition

Processed TCGA Level 3 data on gene expression methylation, copy number alterations, somatic mutations, and microRNA abundance from a total of 3,288 samples of self-reported European ancestry and 17 cancers were downloaded from the Broad Institute Genome Data Analysis Center (GDAC) Firehose on March 18, 2017 using the TCGA2STAT **R** package^30^. Raw genotyping image files (.CEL files from the Affymetrix 6.0 platform) taken from matched normal tissue (i.e., noncancer tissue) were downloaded for each of the 3,288 samples from the National Cancer Institute’s Genomic Data Commons. For all analyses, we only considered tumor samples for which data on gene expression, methylation, copy-number alterations, microRNAs, somatic mutations, and germ-line genetic variation were available (Table 1).

**Table 1:** Sample sizes (*N*) and numbers of non-zero values for TCGA data used to partition variation in gene expression. Cancer sites are abbreviated as: BLCA (bladder urothelial cancer), BRCA (breast invasive carcinoma), CESC (cervical squamous cell Carcinoma), ESCA (esophageal carcinoma), HNSC (head and neck squamous cell carcinoma), KIRC (kidney renal clear cell carcinoma), KIRP (kidney renal papillary cell carcinoma), LGG (brain lower grade glioma), LIHC (liver hepatocellular carcinoma), LUAD (lung adenocarcinoma), PAAD (pancreatic adenocarcinoma), PCPG (pheochromocytoma and paraganglioma), SARC (sarcoma), SKCM (skin cutaneous melanoma), STAD (stomach adenocarcinoma), THCA (thyroid carcinoma), PRAD (prostate adenocarcinoma). RNA-Seq: number of genes with at least one nonzero expression measure; SOM: number of genes with more than one carrier of a somatic mutation; CNA: number of genes with nonzero variability in copy number alterations across patients; METH: number of genes with at least one methylation probe overlapping the TSS ± 1500bp; miRNA: number of microRNAs with at least one nonzero expression measure.

| Cancer | $N$ | RNA-Seq | SOM | CNA | METH | miRNA |
| --- | --- | --- | --- | --- | --- | --- |
| BLCA | 109 | 20,063 | 5,764 | 17,315 | 15,233 | 843 |
| BRCA | 506 | 20,179 | 7,319 | 17,436 | 15,212 | 878 |
| CESC | 136 | 20,016 | 4,685 | 17,310 | 15,157 | 852 |
| ESCA | 113 | 20,223 | 6,269 | 17,450 | 15,242 | 833 |
| HNSC | 245 | 20,134 | 7,360 | 17,377 | 15,235 | 876 |
| KIRC | 228 | 20,150 | 2,463 | 17,401 | 15,264 | 789 |
| KIRP | 95 | 20,008 | 1,147 | 17,287 | 15,208 | 794 |
| LGG | 262 | 20,085 | 1,043 | 17,357 | 15,167 | 833 |
| LIHC | 110 | 19,964 | 2,244 | 17,224 | 15,141 | 817 |
| LUAD | 144 | 19,971 | 6,965 | 17,240 | 15,150 | 832 |
| PAAD | 131 | 19,932 | 3,135 | 17,220 | 15,059 | 792 |
| PCPG | 144 | 19,951 | 246 | 17,249 | 15,160 | 805 |
| PRAD | 132 | 19,983 | 770 | 17,279 | 15,194 | 780 |
| SARC | 210 | 20,176 | 2,343 | 17,402 | 15,150 | 830 |
| SKCM | 320 | 20,172 | 14,297 | 17,406 | 15,225 | 887 |
| STAD | 138 | 20,223 | 9,071 | 17,450 | 15,261 | 826 |
| THCA | 265 | 20,037 | 415 | 17,325 | 15,202 | 851 |

#### Gene expression

Gene expression was measured via RNA-Sequencing on the Illumina Hi-Seq platform and processed using the second TCGA analysis pipeline (RNASeqV2). In this pipeline, per-gene normalized abundance estimates were calculated with the RSEM method^16^. RNA-Seq normalized counts were then log-transformed after adding a constant of 1. In order to correct for any unmeasured confounders or batch effects, we conducted a principal component analysis (PCA) across all genes for each cancer separately as recommended in Leek *et al*. (2010)^15^. For each gene, we regressed the log-transformed RSEM values against the first 5 principal components, and considered the residuals for all subsequent analyses. To ensure that subsequent analyses across genes were comparable, we standardized the residuals to have a variance of 1.

#### Methylation

Methylation was measured on the Illumina Infinium Human Methylation450 Bead-Chip from tumor samples. For each gene, we considered only probes located within 1,500 base pairs of the transcription start site (TSS). The probe with the maximum variance across samples was chosen as the representative measure of promoter methylation for each gene. Probe beta measures, corresponding to the ratio of intensities between methylated and unmethylated alleles, took values between 0 (unmethylated) and 1 (fully methylated) and were transformed to the logit scale prior to our analysis.

#### Somatic mutations

TCGA Level 3 data for somatic mutations are provided as Mutation Annotation Format (MAF) files, which list mutations identified for each patient. To aggregate this information for each individual, `TCGA2STAT` automatically classifies samples as carriers or noncarriers of a nonsynonymous somatic mutation for each gene. We retained this binary coding for all analyses. Unsurprisingly, for most cancers the majority of genes did not have a single carrier of a nonsynonymous somatic mutation. For genes with a single carrier of a nonsynonymous somatic mutation, the somatic mutation component was not included in the model, as this would result in an unstable estimate of its variance component.

#### Copy number alterations

Somatic copy number alterations (CNAs) were called by comparing the Affymetrix 6.0 probe intensities from normal (i.e., non-cancer tissue) compared to probe intensities for cancer tissue. After filtering segments from the Y chromosome, level-3 genome segments provided by TCGA were aggregated to gene-level by `TCGA2STAT` using the `CNTools` Bioconductor package. CNA measures correspond to the log-ratio of copy numbers in the tumor compared to normal samples, where copy number gains and losses correspond to positive and negative values, respectively.

#### Genetic variation

Germ-line genetic variation was available via controlled data access through the Genomic Data Commons (GDC). We downloaded all Affymetrix 6.0 image files (.CEL) files from the GDC derived from normal tissue. For each cancer, we performed genotype calling with the `crlmm` package in `R`. After genotype calling, we performed standard quality control steps including (1) set to missing any genotype called with a quality score less than 0.8; (2) remove samples with a subsequent missing rate greater than 3%; (3) remove markers with a missing rate greater than 3%; (4) remove markers with an minor allele frequency (MAF) less than 1%; and (5) remove any markers with a *P*-value for testing Hardy-Weinberg Equilibrium less than 5 × 10^−8^. For each gene, we considered genetic variants within 1 mega-base of the transcription start site of the gene.

#### microRNA abundance

Data on miRNA abundance were generated on either the Illumina HiSeq 2000 or Illumina Genome Analyzer sequencing machines. Level-3 processed data corresponded to Reads per million microRNA mapped (RPMMM) values. Normalized abundance values were log-transformed after adding a constant of 1 prior to the analysis.

#### Transcription factors

Transcription factors (TFs) play an essential role in regulating gene expression. However, TCGA did not generate data directly measuring transcription factor abundance. In order to integrate TFs into our analyses, we used the expression of the gene (described above) that encodes the TF as a proxy for TF abundance. By crossing the combined lists of TFs provided by Ingenuity Pathway Analysis (IPA; QIAGEN Inc., https://www.qiagenbioinformatics.com/products/ingenuity-pathway-analysis) and the TRRUST database^10^ with the genes for which RNA-Seq expression data were available, we thus characterized the expression of 877 different TFs.

### Linear mixed model analysis and variance partitioning

The goal of our analyses was to partition the variance in gene expression (for each gene) due to promoter methylation, somatic mutations, copy number alterations, microRNA abundance, transcription factor expression, and germ-line genetic variability. To do so, we considered each gene within each cancer type independently, modeling gene expression in a linear mixed model^25^ framework with

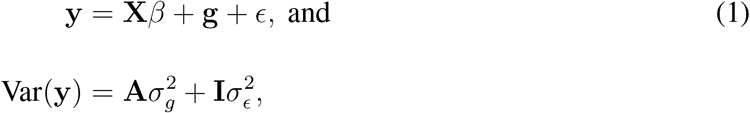

where **y** represents the expression of a fixed gene, **X** is a matrix of fixed effects, **g** is a vector of the total genetic effects across samples, and 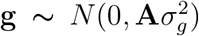 where **A** is obtained from the germ-line genetic data to represent a genetic relationship matrix between individuals, as in Yang *et al*. (2011)^33^. The columns of **X** correspond to the non-genetic factors contributing to variation in expression, i.e., somatic mutations, CNAs, methylation, miRNAs, and transcription factors.

This linear mixed model represents a powerful and flexible statistical framework that is based on well-characterized theoretical properties and facilitates both the joint quantification of multiple sources of variation as well as a comparison of their relative contributions.

#### Variable selection for TFs and miRNAs

TFs and miRNAs can each potentially target multiple genes, and a substantial body of work has emerged to develop methods and databases to predict these TF-target and miRNA-target pairs, for example using text-mining^10^ or bioinformatics-driven approaches^32^. However, as these regulatory interactions are complex and knowledge of the biological processes driving them is far from complete, identifying a definitive mapping of TFs and miRNAs to genes from the large lists of available predicted targets is not straightforward.

For both TFs and miRNAs, several hundred expression measures were available, meaning that the inclusion of all measured TFs and miRNAs in each of our gene-level models was not possible. As such, rather than making use of databases of predicted TF- and miRNA-target pairs to reduce the dimensionality of our data, we adopted an alternative approach. Specifically, we implemented a sparse principal component analysis (sPCA)^27,35^ of TF and miRNA expression using the `mixOmics` packge in R. This analysis serves two purposes: (1) dimension reduction to preserve degrees of freedom for the estimation of the linear mixed model; and (2) enhanced interpretability of our results by identifying the top TFs and miRNAs that contributed to overall variation in gene expression. The number of nonzero loadings for each sPC was set to 10. Because sPCs may be correlated (in contrast to standard PCs which are always orthogonal), we selected the top 5 sPCs for TF and miRNA expression that were uncorrelated with one another (absolute Pearson correlation < 0.3). The TFs and miRNAs with non-zero loadings in at least one of the sPCs thus correspond to those that contribute the most to variation in overall TF and microRNA expression, respectively, for each cancer site.

We followed this approach for selecting sPCs for every cancer, with the exception of brain lower grade glioma (LGG). We noticed that the top TF sPCs for LGG were effectively proxies for the concurrent hemizygous deletions of the 1p and 19q regions, common to specific sub-types of LGG^29^. For this special case, we re-selected the top TF sPCs that were uncorrelated with the CNA data and that specifically captured the hemizygous deletions.

#### Model fitting

In order to fit the linear mixed model, we implemented a restricted maximum likelihood (REML) procedure as follows. We obtained standardized residuals after regressing gene-level RSEM values against the first 5 transcriptome-wide PCs (as described above). These standardized residuals were then regressed against the following fixed effects: (1) promoter methylation levels; (2) somatic mutation carrier status; (3) CNA values; (4) the top 5 uncorrelated sPCs representing variation in miRNA levels; and (5) the top 5 uncorrelated sPCs representing variation in TF levels. For every gene, the residuals from this model were then input to the GCTA (version 1.26.0) software^33^ along with all germ-line genetic polymorphisms measured within 1 mega-base of the TSS for each gene, similar to the approach in Gusev *et al*. (2016)^9^, to estimate the component of variance due to cis-acting genetic variation 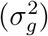. Variance components for the fixed effects were estimated as 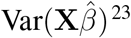^23^, and variance components for the random genetic effect were estimated with REML as implemented in GCTA. In order to avoid confounding by potential population structure, we restricted our analyses to the largest population, individuals of self-reported European ancestry.

### Promoter reporter assay for *PTPN14*

A 1232bp segment of the human PTPN14 promoter (−640bp to +589bp from the transcriptional start site) was cloned into the pEZX-PG02.1 GLuc-ON^™^ promoter reporter construct (cat# LF061, GeneCopia, Rockville, MD). MDA-MB-231 cells were transfected with 0.5 *μg* of DNA composed of 0.25 *μg* of the PTPN14 promoter-reporter construct and 0.25 *μg* of mammalian expression vectors encoding the human FOXOA1, GATA3, or empty vector control. At 24 hours post-transfection, media was collected and luciferase activity was measured following the manufacturer’s protocol (Luciferase assay; GeneCopoeia).

## Results

### *EDGE in TCGA* Shiny App

In order to facilitate rapid exploration of the results from our partitioning of variance in transcriptome-wide gene expression across 17 different cancers, we developed an interactive `R` Shiny App called *EDGE in TCGA* for visualization, browsing of genes and cancers, and identification of important TFs and miRNAs. Variance components from every gene were organized into sortable tables for easy within cancer gene-based lookups^22^ and inter-active plots, including heatmaps^7,8^ and lollipop plots^31^, for pan-cancer gene-based explorations. For the TF and microRNA sPCs, we summarized the contribution of TFs (and microRNAs) by multiplying the corresponding estimated coefficients 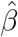 from the linear mixed model with the loadings from the sPC decomposition. The *EDGE in TCGA* tool is freely available at http://ls-shiny-prod.uwm.edu/edge_in_tcga.

### Pan-cancer trends in gene expression drivers

We used the *EDGE in TCGA* tool to explore pancancer trends in the drivers of gene expression. Across the transcriptome and for all cancers that we analyzed, CNAs represented the most consistent driver of gene expression, with somatic mutations and germ-line genetic polymorphisms influencing much smaller numbers of genes (Table 2), consistent with previous reports of the relative importance of aneuploidy versus somatic mutations or germ-line polymorphisms^6,17^. The number of genes with notable (i.e., > 10% estimated variance component) genetic effects appear to be similar across cancers with the exception of PRAD and KIRP, which have the highest number of genes with germ-line genetic drivers expression. The effects of miRNAs also appear to be similar across cancers, with LUAD and LIHC as clear outliers with a large number of genes affected by miRNA variation. However, the relative influence of methylation, CNAs, and TFs varied considerably across cancers; when comparing the variance due to each of these effects across cancers, we observed distinct clustering of cancers for some genes.

**Table 2:** The number of genes with over 10% of the variance in their expression due to the effect of: MUT (somatic mutations), CNA (copy number alterations), METH (methylation), miRNA (microRNA), TF (transcription factors), and GEN (cis-genetic polymorphisms).

| Cancer | MUT | CNA | METH | miRNA | TF | GEN |
| --- | --- | --- | --- | --- | --- | --- |
| BLCA | 29 | 8020 | 1346 | 3193 | 2119 | 857 |
| BRCA | 3 | 7838 | 946 | 352 | 846 | 161 |
| CESC | 33 | 7866 | 1455 | 1537 | 3398 | 700 |
| ESCA | 18 | 8434 | 1582 | 1436 | 2606 | 790 |
| HNSC | 6 | 7858 | 936 | 298 | 1258 | 264 |
| KIRC | 5 | 5545 | 614 | 1244 | 639 | 475 |
| KIRP | 21 | 7318 | 1379 | 1703 | 4983 | 1109 |
| LGG | 3 | 5652 | 467 | 687 | 5458 | 438 |
| LIHC | 12 | 7094 | 1356 | 3097 | 4801 | 911 |
| LUAD | 21 | 7868 | 1239 | 4075 | 3935 | 608 |
| PAAD | 7 | 7403 | 855 | 1569 | 2312 | 874 |
| PCPG | 2 | 6721 | 424 | 980 | 3289 | 779 |
| PRAD | 4 | 5412 | 1122 | 977 | 2246 | 1022 |
| SARC | 4 | 8185 | 927 | 334 | 1875 | 377 |
| SKCM | 6 | 7539 | 1358 | 947 | 1896 | 135 |
| STAD | 25 | 7609 | 1450 | 1482 | 2837 | 672 |
| THCA | 2 | 2261 | 391 | 2457 | 2623 | 617 |

As one example, Figure 1 illustrates the relative molecular variance components for genes encoding components of the p53-DNA repair pathway, a major oncogenic pathway responsible for maintaining the fidelity of DNA replication and cell division. Although germ-line genetic mutations and promoter methylation have relatively weak roles in driving the expression of genes in this pathway, CNAs tend to have much larger effects in this pathway across cancer types. Interestingly, two genes in this pathway (BRCA1, BRCA2) have large variance components related to TF expression in a subset of four cancers (LGG, SARC, LUAD, SKCM). By weighting the non-zero sPC loadings with their respective coefficients in the linear mixed model in Equation (1), we can evaluate the relative importance of specific TFs to the expression of the target gene in these cancers. Figure 2 shows the “TF contribution” tab in the *EDGE in TCGA* tool for LGG and SKCM where we observe several similarities among the set of TFs acting as moderate drivers of the expression of BRCA2; for instance, FOXM1, RAD51, and MYLBL2 are among the largest positively associated TFs in both cancers. However, the TFs corresponding to the largest positive association with the expression of BRCA2 in LGG and SKCM are E2F1 and UHRF1, respectively. This suggests that although BRCA2 expression in these two cancers appear to be regulated by similar TF programs, there are also unique differences between the two cancer types.

**Figure 1:**
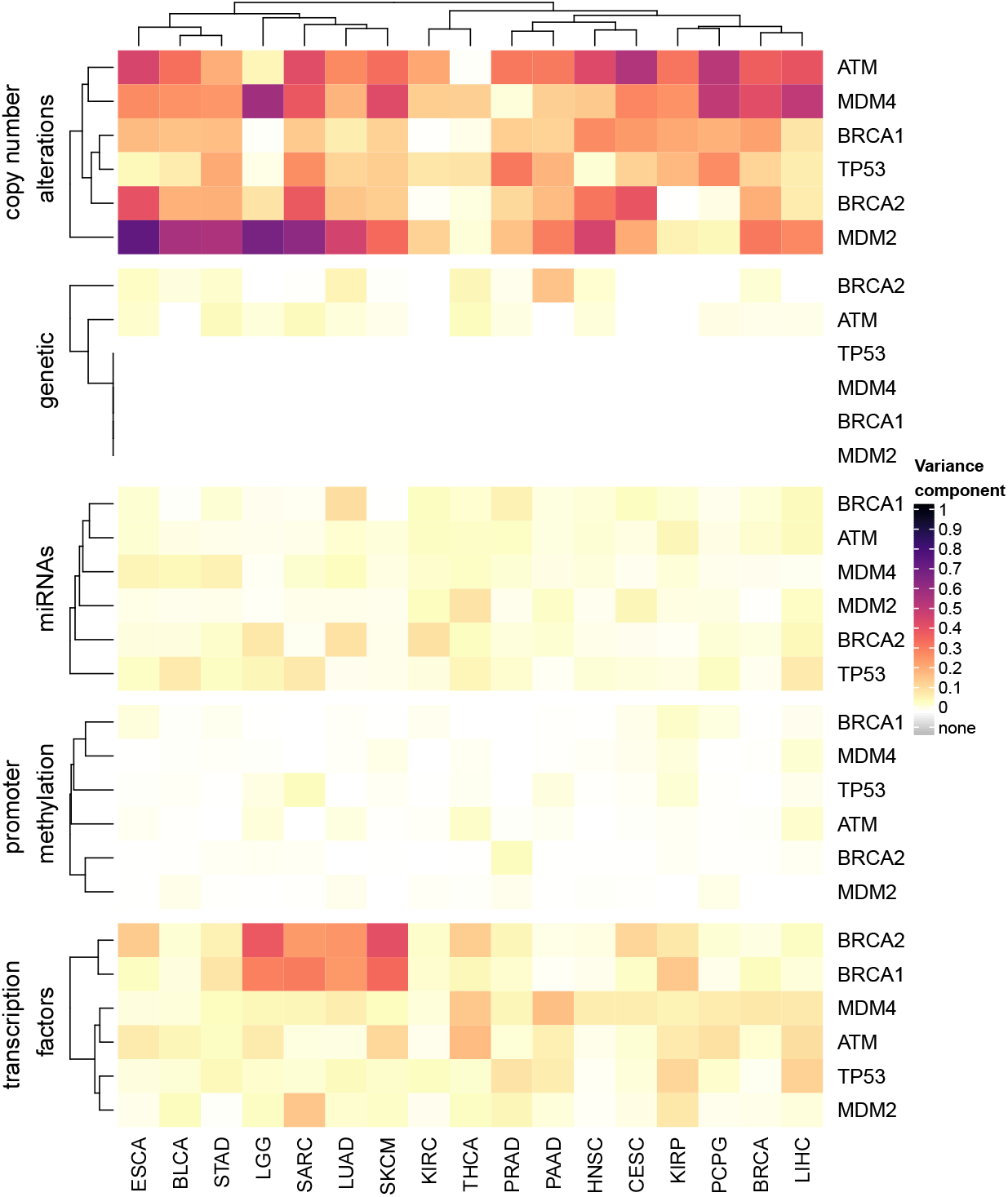
Variance component estimates for CNAs, germ-line genetic polymorphims, promoter methylation, miRNA abundance, and TF expression for genes in the p53-DNA repair pathway (BRCA1, BRCA2, ATM, MDM2, MDM4, TP53).

**Figure 2:**
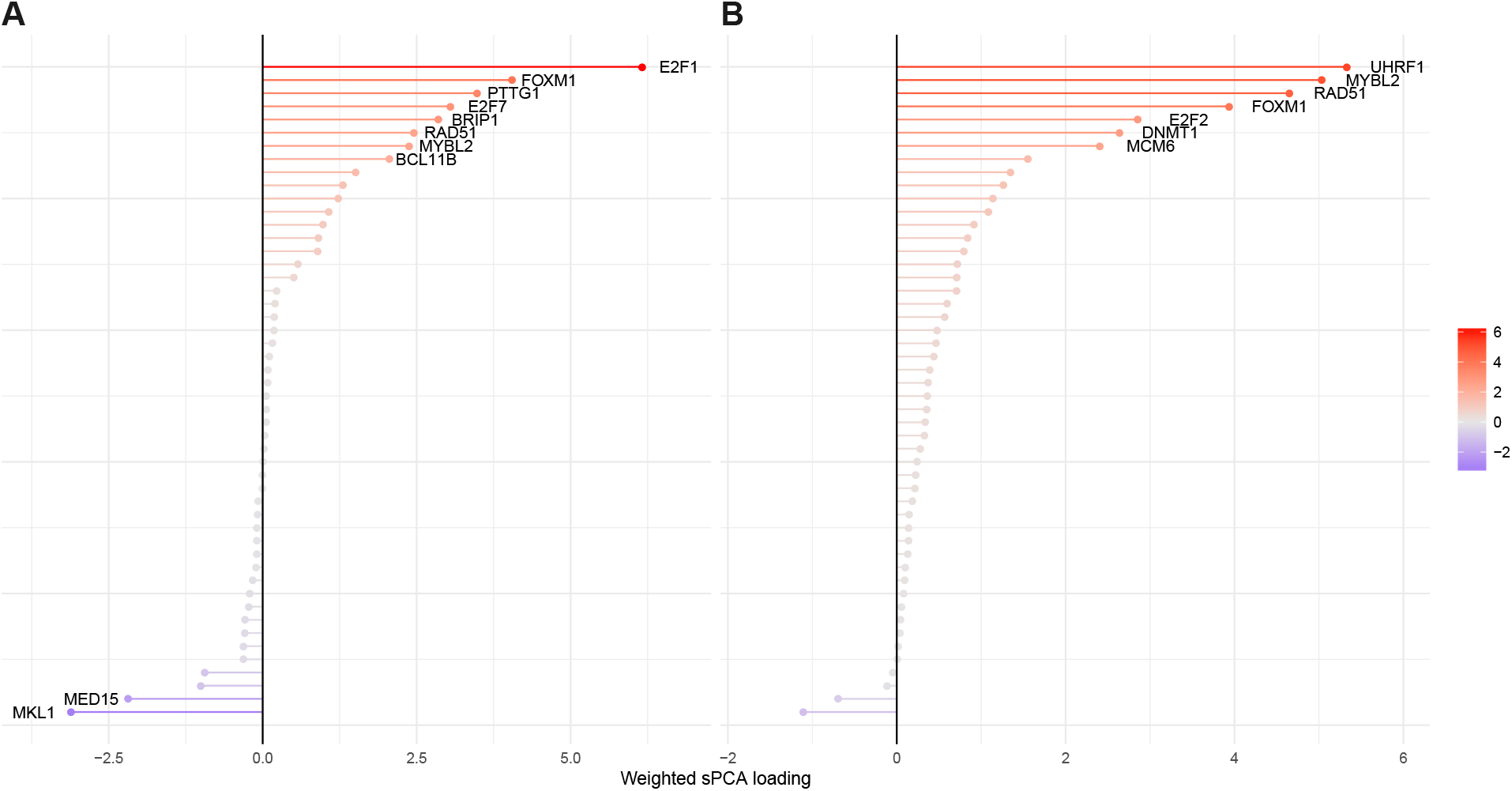
Lollipop plots of transcription factors associated with BRCA2 gene expression in LGG (A) and SKCM (B). Non-zero loadings from the first five uncorrelated sparse principal components, weighted by their respective coefficients from the linear mixed model in Equation (1) in LGG and SKCM. Weighted loadings are plotted from most strongly negative (blue) to most strongly positive (red).

It may also be of interest to examine the pan-cancer trends in genetic and epigenetic drivers of expression in specific genes. For example, MYC is a known oncogene that encodes a protein involved in many cellular functions, including cell cycle progression and DNA replication. So-called C-class tumors^3^ dominated by multiple recurrent chromosomal gains and losses were found to be characterized in part by MYC-driven proliferation. Across cancer sites, the dominant drivers of MYC expression varied widely among cancers (Figure 3). Cancer sites largely grouped into one with a large miRNA component (LUAD), those with large CNA drivers (BLCA, PAAD), those with large TF drivers (PCPG, LGG, THCA), and those with both CNA and TF drivers of expression variation (LIHC, ESCA, STAD). For a large set of cancer sites (PRAD, KIRP, SARC, KIRC, SKCM, HNSC, BRCA, CESC), the residual variance component was predominant.

**Figure 3:**
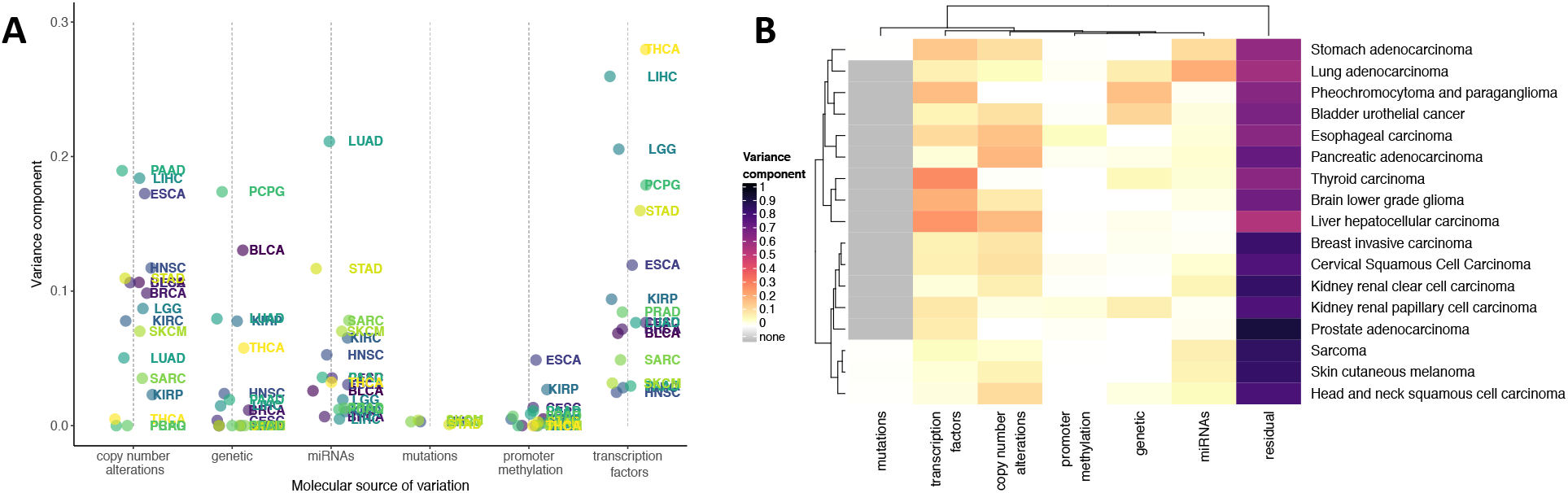
(A) Variance component estimates for MYC expression for each molecular source of variation in each of the 17 cancers. (B) Heatmap of the variance component estimates for MYC expression in each of the 17 cancers, with the estimated heritability, mean logit-transformed methylation *β* values across samples, percent samples with somatic mutations, mean normalized CNA values across samples, and mean log-normalized RNA-seq expression across samples for each cancer.

### *PTPN14* promoter reporter assay

Beyond an exploratory pan-cancer analysis of global patterns, the estimated variance components can also be used to investigate candidates within specific cancer types. For instance, PTPN14 is a protein phosphatase that has been implicated in breast cancer risk^34^ and acts as a tumor suppressor in breast cancer^1^ and other malignancies^20,28^, yet the mechanisms that regulate PTPN14 expression are largely unknown. Using the *EDGE in TCGA* tool, we explored the mechanisms regulating PTPN14 expression in breast cancer (BRCA). In BRCA, a large amount of variance in PTPN14 expression was explained by CNA and TF, suggesting that multiple drivers may underlie the dysregulation of PTPN14 expression in breast cancer. To explore how TFs potentially regulate PTPN14 expression, we examined the existing ENCODE database to identify transcriptional regulators that bind to the PTPN14 promoter in T47D breast cancer cells, which revealed binding sites for GATA3 and FOXA1 (Figure 4A). Likewise, the “TF contribution” tab in the *EDGE in TCGA* tool revealed that GATA3 and FOXA1 expression were strongly correlated with PTPN14 expression in BRCA and suggested an inverse relationship (Figure 4B). The ability of GATA3 and FOXA1 to repress the PTPN14 promoter was tested using a PTPN14 promoter-luciferase reporter assay. Compared with the empty vector control group, the PTPN14 promoter activity was significantly downregulated by co-expression of GATA3 (~5-fold, *P* < 0.001) and FOXA1 (~3-fold, *P* < 0.001) (Figure 4C), suggesting that GATA3 and FOXA1 indeed repress the PTPN14 promoter. Collectively, these data demonstrate the utility of the *EDGE in TCGA* tool to identify mechanisms that are biologically relevant and can be tested at the molecular level.

**Figure 4:**
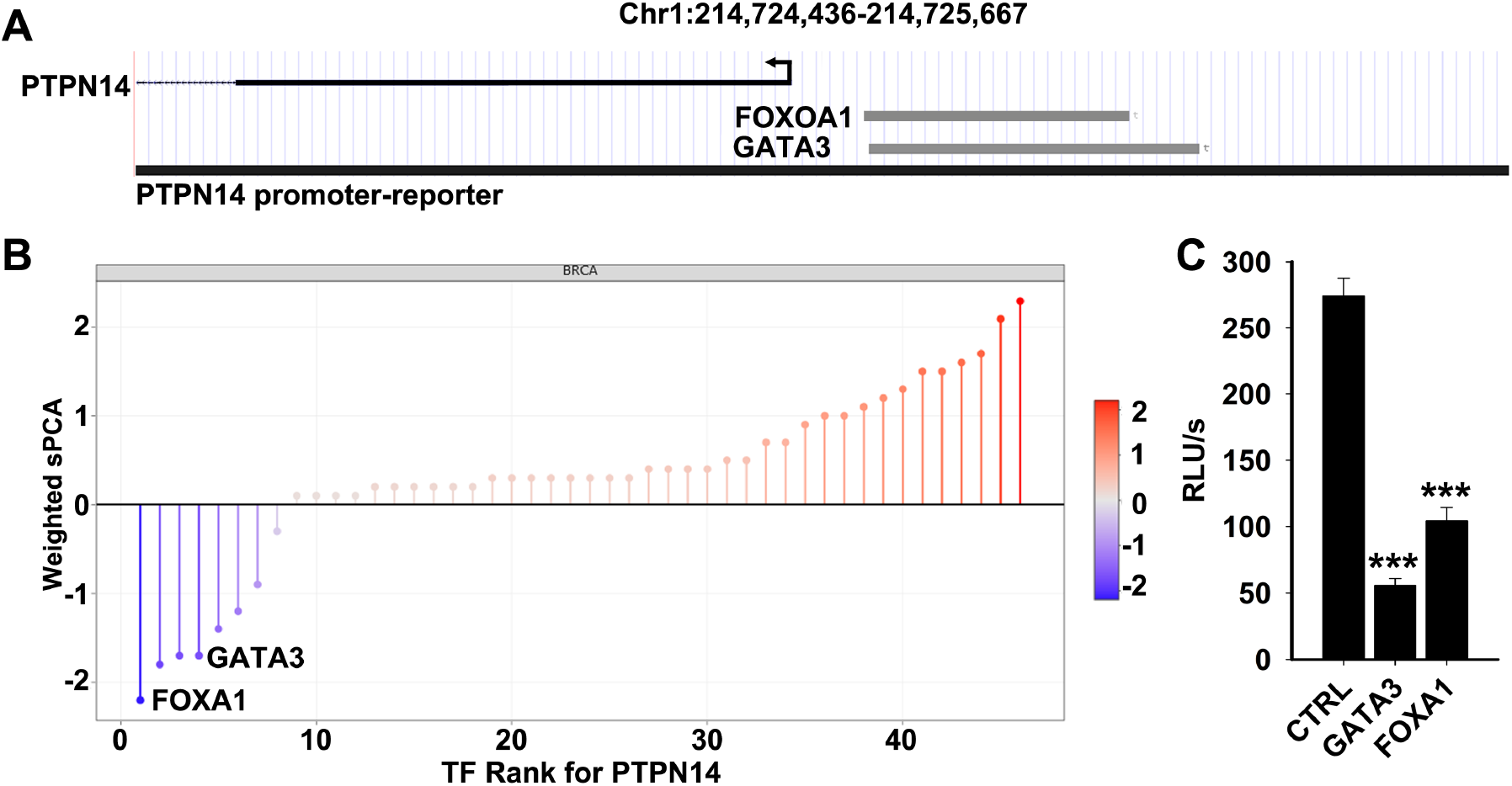
(A) Transcription factor tracks from the UCSC genome browser of FOXA1 and GATA3 binding at the promoter of PTNP14 in ENCODE breast cancer cell line T47D. (B) TF zoom plot of the relative importance of different transcription factors on the expression of PTPN14 in breast cancer tumor samples. The vertical axis shows the sparse PCA loadings scaled by the effects of each sPC. Transcription factors are ranked from most strongly down-regulating gene expression (bottom left) to most strongly up-regulating gene expression(top right). (C) Relative expression of PTNP14 with FOXA1 and GATA3 introduced, versus controls.

## Discussion

Data from the TCGA project have been used in a multitude of contexts to explore the molecular basis of cancer. The genome-wide results from our agnostic, integrative analysis of the molecular drivers of gene expression in TCGA tumor samples provide a new way of exploring the TCGA data. Browseable results in the *EDGE in TCGA* Shiny App can be used to generate hypotheses and offer unanticipated insights into the molecular basis of a number of different cancers.

As an example, we prioritized the transcription factors that are likely to govern the expression of *PTPN14* in breast cancer cells. Subsequent experiments with promoter reporter assays confirmed that FOXA1 and GATA3 (implicated as important TFs in our analysis) regulate the expression of *PTPN14* in a breast cancer cell-line. In addition to the identification of important TFs, our *EDGE in TCGA* Shiny App can be used similarly to identify important miRNAs, methylation sites, somatic mutations, and copy-number alterations that regulate gene expression in 17 different cancers.

Though the *EDGE in TCGA* Shiny App provides a powerful tool for exploring the drivers of gene expression in the TCGA data, it comes with certain caveats that should be carefully considered when interpreting the results. First, we explicitly did not undertake statistical hypothesis testing and do not provide *P*-values for any of our estimated effects. This was done for two reasons: (a) a well-known drawback of linear mixed models is the instability of *P*-values for testing the significance of random effects^25^; and (b) rather than making inferences about gene expression generally, our results are best thought of as a useful summary of the TCGA data to be used for future, hypothesis-driven exploration. Second, we provide a tool for comparing the relative importance of fixed effects (MUT, CNA, METH, miRNA, TF) and random effects (GEN) on the expression of a specific gene. We do not provide any measures of absolute importance and the results should not be interpreted in this way. For example, the effects shown on the TF zoom plots (Figure 2) do not represent effect sizes that estimate an absolute quantity; a value of −3 on this plot means that the associated TF down-regulates expression of the gene 3 times more than a TF with a value of −1.

As more large-scale omics data continue to be generated (for example, through the National Heart, Lung, and Blood’s (NHLBI) Trans-Omics for Precision Medicine (TOPMed) program), there will be renewed interest in “integrative” analyses that bring together data on germ-line genetics, gene expression, methylation, proteomics, and metabolomics. We introduce a statistical framework for partitioning the variation in gene expression due to a variety of molecular traits. This partitioning of variation in gene expression has been performed extensively in the context of germ-line genetic variation^5,9^. We have extended this framework to include other important drivers of gene expression in tumor samples, such as somatic mutations, TFs, miRNAs, CNAs, and methylation. Though our results are specific to the 17 cancers that were included here, the analytic structure is applicable to any phenotype for which multiple matched omics data may be generated.

## Acknowledgements

We thank the developers at RStudio for the Shiny App platform as well as the developers of all the supporting `R` packages. The results shown here are in whole based upon data generated by the TCGA Research Network: http://cancergenome.nih.gov.

## Availability

All TCGA Open Access tier data are available at the Broad Institute GDAC Firehose and were downloaded using the TCGA2STAT **R** package. TCGA Controlled Access tier data are available via controlled access through the Genomic Data Commons (GDC). `R` scripts used to download, format, and analyze the data and produce the interactive R/Shiny web app have been made available on GitHub at https://github.com/andreamrau/EDGE-in-TCGA.

## Correspondence

Correspondence and requests for materials should be addressed to PLA.

